# Tuning the gate and the gear: The LRRC26 (γ1) subunit modulates intrinsic gating and voltage-sensor coupling of the BK channel

**DOI:** 10.1101/2025.09.04.674290

**Authors:** Felipe Echeverria, Antonio Peña-Pichicoi, Miguel Fernandez, Willy Carrasquel-Ursulaez, Juan P. Castillo, Osvaldo Alvarez, Ramon Latorre

## Abstract

Association of auxiliary subunits (β1-4 and γ1-4) with the pore-forming α subunit of the calcium- and voltage-activated potassium (BK) channel provides functional diversity. γ1 promotes a significant leftward shift of the voltage activation curve, ensuring the adequate functioning of secretory glands, allowing the BK channel to release K^+^ at the cell’s resting Ca^2+^ concentration. Given its physiological importance, it is crucial to elucidate the mechanisms of γ1 action. However, structural and functional studies have yielded conflicting conclusions regarding the modulation of BK channels by γ1. Here, using macroscopic, single-channel, and gating current measurements, we demonstrate that at zero mV γ1 increases 92-fold the equilibrium constant that defines the closed-open transition by destabilizing the channel’s closed configuration and enhancing the coupling between the voltage sensor and the pore domain, without affecting voltage-sensor activation. These results suggest that γ1 not only causes an increase in the energetic coupling between the voltage sensors and the pore but mainly enhances the channel opening reaction.

**Teaser:** The γ1 subunit favors the BK channel pore opening by destabilizing its closed configuration.

## Introduction

Large-conductance calcium- and voltage-activated potassium channels (BK, MaxiK, Slo1, or KCa_1.1_) are transmembrane proteins that transport with high selectivity K^+^ over other ions (*1*) with a remarkably high unitary conductance compared to other K^+^ channels (*2, 3*). The membrane potential and a wide range of intracellular Ca2+ concentrations independently modulate its activity (*4, 5*) allowing BK channels to have a pivotal role in many physiological processes such as action potential transmission and neurotransmitter release in nervous system (*6–8*) or vascular tone and contractility in smooth muscle (*9, 10*)

BK channels are homotetramers where each pore-forming (α) subunit contains seven transmembrane segments (S0-S6) and an intracellular motif that mediates activation by intracellular Ca^2+.^ (*8*). They are modular proteins where every subunit contains a voltage-sensing domain, VSD (S0-S4), a pore domain, PD (S5-P-S6), and four carboxy-terminal domains (CTD) that associate to form a gating ring containing the Ca^2+^-binding sites. (*11*). Even though a single gene codifies the BK α subunit – *Slowpoke, Kcnma1* – presents a great variety of channel phenotypes due to alternative splicing of pre-mRNA (*12–14*), metabolic regulation (*15*), and association with auxiliary subunits. The modulation of the BK channel, mediated by β-subunits (β1-β4), has been extensively studied due to its crucial role in the physiological functioning of tissues and organs, including the nervous system, smooth muscle, and kidney. (*8, 16, 17*). The discovery of the γ-subunits (γ1-γ4) increased the diversity of BK channels (*18, 19*) and, more recently, the finding of LINGO1. Coexpression of the BK channel α-subunit with LINGO results in rapidly inactivating K^+^ currents (*20*).

Since the first electrophysiological characterization of BK channels, it was apparent that there is a wide range of BK channel Ca^2+^ sensitivities (*21*). BK channels with high Ca^2+^ sensitivity were identified in different cell types, such as mouse and rat inner hair cells (*22*), human glioma cells (*23*), mammalian salivary, lacrimal, and secretory epithelial glands, (*24–27*) and colon (*28*). Aldrich’s group showed that the high Ca^2+^ sensitivity in LNCaP prostate cancer cells (*29*) was due to the presence of a BK auxiliary subunit, a leucine-rich repeat (LRR)-containing protein 26 (LRRC26; γ1), which leftward-shifted BK voltage-activation curves around 140 mV compared to BK(α) alone in the absence of intracellular Ca^2+^ (*18, 19*). The family of LRR is composed of three other paralogs γ2-γ4 (LRRC52, LRRC55, and LRRC38, respectively), which also modulate BK channels with a stoichiometry α:γ up to 1:1 (*18, 19, 30*). The four γ subunits share a common structure: an N-terminal signal peptide, an extracellular LRR domain, a single transmembrane segment, and a short C-terminal containing a cluster of arginine residues (*31, 32*). γ2-γ4 subunits are also able to modulate the BK channel, albeit with less potency than γ1 (*19*) due to sequence differences, particularly in TM and C-terminal (*33*).

The BK channel associated with γ1 (BK(*α*+γ1)) complex appears to be expressed exclusively in secretory epithelial cells, where it plays a crucial role in K^+^ secretion. The large leftward shift in the voltage axis promoted by the γ1 subunit at cell resting Ca^2+^ concentrations, (*19*) becomes crucial in the adequate functioning of secretory cells contained in the lacrimal gland, parotid gland, and colon (*27*). In cells of the lacrimal gland and parotid gland coming from γ1 knockout mice, BK channels behave as channels containing only the α-subunit (*27*). γ1 has also been found in breast cancer cells and is a requisite to promote the cell malignant phenotype (*34*). Compared to β-subunits, which incrementally increase the probability of opening, a single γ1 subunit can induce a complete gating shift (*30, 35–37*).

Recently, three independent groups have solved the structure of the BK(*α*+γ1) channel complex (*38–40*). Two of these structures display the whole γ1 subunit (*38, 40*), and the third structure contained the γ1 transmembrane segment only (*39*). These structural studies confirmed that the complex has a stoichiometry of α:γ1 = 1:1 as previously inferred using electrophysiological and fluorescent techniques (*30, 35*). In the BK(*α*+γ1) channel, the γ1 transmembrane segment adopts a kink that interacts with S0-S3 and appears to be a critical structural element for the modulatory effects of the γ1 subunit (*41*). The arginine cluster located at the C-terminal of the γ1 is also crucial in determining the BK gating changes. By making chimeras in which the different domains of the different γ subunits were exchanged, Li et al. (*31*) determined that the differential modulation of BK channels induced by the different γ subunits was determined by their transmembrane domain and the positively charged C-terminal. On the other hand, the LRR domain regulates subunit trafficking, and its deletion produces BK channel in which the all-or-none effect is absent (*42*).

The mechanism by which this positively charged cluster modulates the BK channel function is under debate. Analysis of the electrophysiological results using the Horrigan-Aldrich (HA) allosteric model (*43*) (**Fig. S1A**) suggested that the presence of the γ1 subunit mainly affects the coupling between the VSD and the PD (*18*). However, a recent structural study of the γ1-containing BK channel, suggests that γ1 stabilizes the active conformation of voltage and Ca^2+^ sensors, as well as the strength of interaction between these two sensors (*40*). In particular, the intracellular interaction of the γ1 arginine cluster with the RCK1 lobe suggested that the γ1 modulates BK Ca ^2+^ and voltage activation (*40*). On the other hand, Redhardt et al. (*39*) hypothesized that the arginine cluster introduces an electrostatic positive bias on the internal aspect of the BK channel, stabilizing the active conformation of the VSD. A different view is put forward by Kallure et al. (*38*) which suggested that the arginine cluster present in the γ1 C-terminal domain plays a role in the association between the γ1 and α subunit. They pointed out that positively charged residues at the end of transmembrane segments interact with phospholipids, helping the membrane integration of the γ1.

Due to the different interpretations of the structural data about the γ1 mechanism of action and lacking functional conclusive evidence, this study aims to clarify γ1 modulation using an electrophysiological approach along with the HA model, which includes measuring ionic and gating currents in multiple Ca^2+^ conditions to determine which parameters in the allosteric gating model are affected by the γ1 subunit. Our data indicates that while the γ1 subunit increases the coupling between the VSD and the PD, its main effect is increasing the PD equilibrium constant (*L*; **Fig. S1**) by destabilizing the closed conformation.

## Results

### The γ1 subunit does not alter the resting-active equilibrium of the voltage sensor

We expressed the BK α subunit in the presence and absence of γ1 in *Xenopus laevis* oocytes. In both conditions, we observed robust K^+^ currents in the absence of Ca^2+^ (**Fig. 1A-B**). The plot of the normalized tail currents vs. voltage (G(V)) curves shows that, as previously found (*18, 41*) in the absence of Ca^2+^, γ1 produces a considerable leftward shift that amounts to −137 mV (**Fig. 1C**, see **Table S1**). This value is like those found previously using PC3 cells (−136 mV) or HEK cells (−147 mV) (*18*) and in *Xenopus* oocytes (−120 mV)(*36*). Fig. 1B also shows that the deactivation kinetics becomes slower in BK(α+γ1) compared to BK(α) channels.

**Figure 1.**
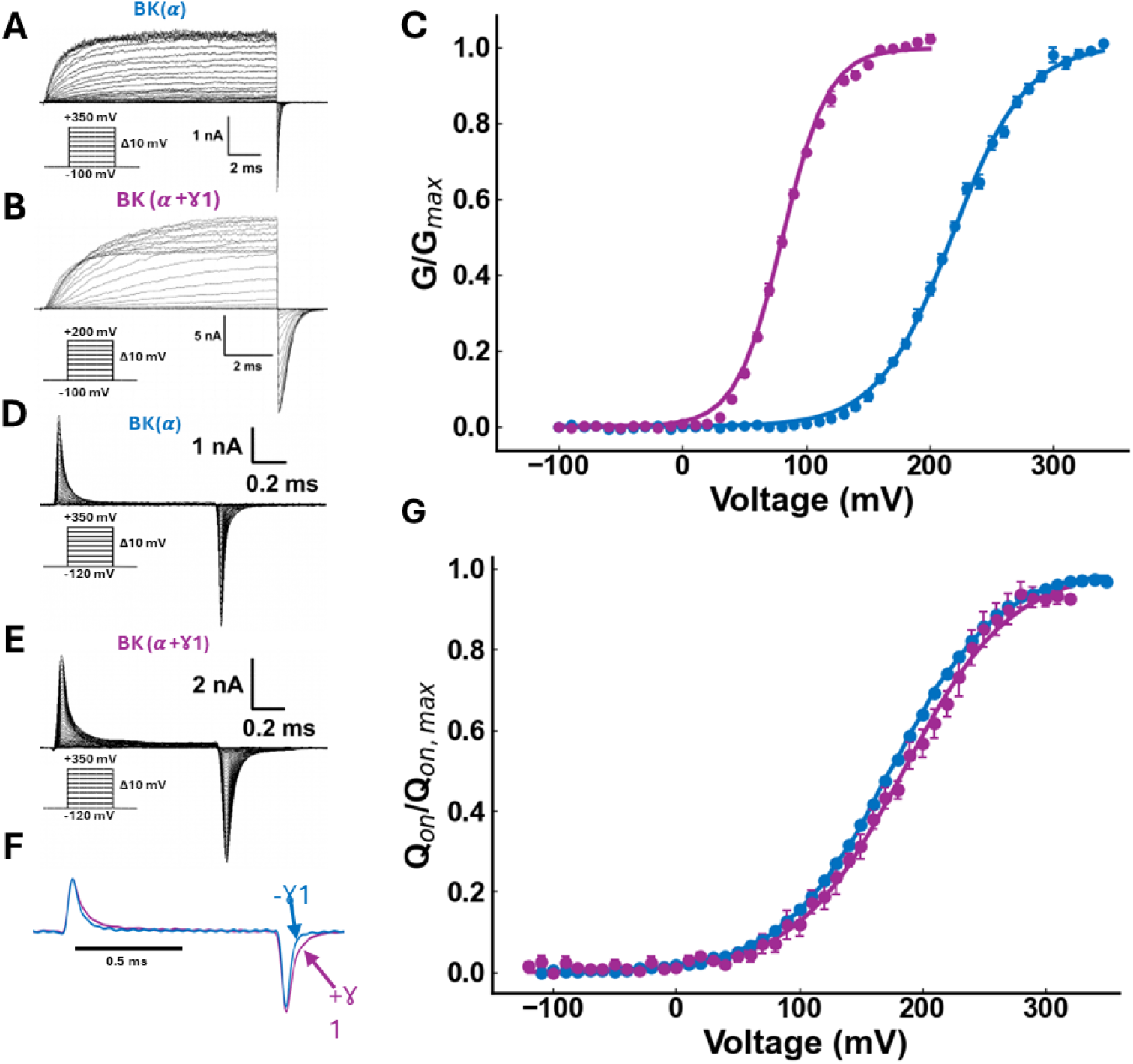
γ1 subunit increases BK channel open probability without affecting the resting-active equilibrium of the voltage sensor. Potassium currents recorded in inside-out membrane patches containing (**A**) BK(*α*) channels or (**B**) BK(*α*+γ1) channels. Currents were recorded under symmetric 110 mM K^+^ conditions and zero µM free internal Ca^2+^. Inset: Voltage protocol used. (**C**) Normalized conductance versus voltage plots obtained from tail currents of BK(*α*) (blue, N = 9) and BK(*α*+γ1) (purple, N = 8). Data were represented as mean ± SEM and fitted using the relation G/Gmax = 1/[1+exp(−z_G_F(V-V_0.5_)/RT)] where z_G_ is the apparent gating valence, V the voltage, and V_0.5_ the half-voltage. BK(*α*): z_G_ = 0.85 ± 0.02 and V_0.5_ = 218.3 ± 0.8 mV. BK(*α*+γ1): z_G_ = 1.37 ± 0.02 y V_0.5_ = 80.9 ± 0.3 mV. (**D**) Gating currents in inside-out membrane patches containing BK(*α*) channels and (**E**) BK(*α*+γ1) channels, both recorded under conditions of zero µM free internal calcium. Inset: Voltage protocol used. (**F**) Normalized gating currents evoked by 1-ms pulses to 210 mV for BK(*α*) (blue) and BK(*α*+ γ1) (purple). Holding voltage −100 mV. A single exponential function was needed to fit the ON-gating currents of BK(*α*), and a double exponential function was needed to fit the ON-gating currents elicited by BK(*α* + γ1). (**G**) Q(V) data obtained from the area under the fastest exponential function were plotted vs. voltage and fitted using a two-state model: *Q*_*on*_*/Q*_*on-max*_ = 1/[1+exp(−z_Q_F(V – V_H_)/RT)]. BK(*α*): z_Q_ = 0.59 ± 0.04 and V_H_ = 178 ± 0.4 mV (N = 4), while for BK(*α*+γ1): z_δ_ = 0.58 ± 0.04 and V_H_ = 185 ± 1.5 mV (N = 5).

We next asked the question: Does the γ1 subunit modify, and to what extent, the voltage sensor’s resting-active (R-A) equilibrium? We recall here that the equilibrium constant (*J*) defining the R-A equilibrium is given by (*43*):

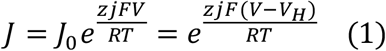

where *J*_*0*_ is the equilibrium constant at zero mV, *z*_*J*_ defines the voltage dependence of the voltage sensor activation, *V* is the applied voltage, *V*_*H*_ is the half activation voltage, and *F, R*, and *T* have their usual meanings. The normalized total gating charge displaced during activation (*Q*_*on*_*/Q*_*on*.*max*_) relates to the equilibrium constant *J* according to the relationship,

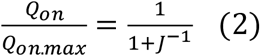

We characterize the effect of the γ1 subunit on the R-A equilibrium by measuring gating currents in the absence and the presence of γ1 (**Fig. 1D-E**). It is apparent that the kinetics of the ON-gating current in BK(α+γ1) has two exponential components, unlike BK(α) channels, which, as found previously (*43–45*) can be approximated to a single exponential function (**Fig. 1F - Fig. S2**). Moreover, the presence of γ1 significantly slows down the OFF-gating current (**Fig. 1F**).

Integration over time of the fast component of the ON-gating currents shows that the presence of γ1 do not alter significantly neither the parameter *z*_*J*_ (0.58 ± 0.04 for BK(α+γ1); 0.59 ± 0.04 for BK(α)) nor *V*_*h*_ (184.7 ± 1.5 mV for BK(α+γ1); 178.4 ± 0.4 mV for BK(α)) obtained when fitting the Q(V) data to Eq. 2 (**Fig. 1G**, see **Table S1**). Thus, this result shows that γ1 modifies the voltage sensor’s R-A equilibrium marginally, promoting a 7-mV rightward shift of the Q(V) curve. A comparison of the G(V) and Q(V) curves in the presence of γ1 reveals that 90% of the maximum conductance occurs when only ~20% of the voltage sensors are active. I.e., γ1 allows the BK channel to open when fewer voltage sensors are activated. Since the fast component of the ON-gating current reflects the voltage sensor movement when the channels are in the closed state, Fig. 1G allows us to conclude that the γ1 subunit promotes only a minor change in the equilibrium constant for voltage sensor activation, *J*_*0*_ *= exp(−z*_*J*_*FV*_*h*_*/RT*) (**Eq. 1**, see **Table S1**).

### The γ1 subunit modulates the open-closed equilibrium

In the absence of Ca^2+^, the allosteric model (**Fig. S1A**) predicts that the channel open probability (*P*_*o*_) is given by (*43*):

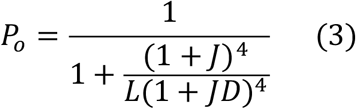

where 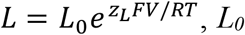 is the closed-to-open equilibrium constant at zero mV, *z*_*L*_ is the voltage dependence for channel opening, and *D* is the allosteric factor that couples the voltage sensor with the pore opening.

To have a complete description of the nature of the effect of γ1 on *P*_*o*_, in the absence of Ca^2+^, we need to determine the parameters *L* and *D* in Eq. 3. Since the equilibrium constant *J* (**Eq. 1**) vanishes for very negative voltages, **Eq. 3** tends *Po ≈ L(V*). Therefore, we can obtain *L*_*0*_ and *z*_*L*_ by plotting ln(*Po*) versus voltage when all the voltage sensors are at rest. This plot on a semilogarithmic scale is a straight line with a 0-volt intercept *L*_*o*_, and the slope is *z*_*L*_*F/RT*. To get *Po*, we detect single-channel activity at negative voltages in membrane patches containing a known number of channels (*N*). Fig. 2A-B shows single-channel records obtained at two different voltages in the absence and presence of the γ1 subunit, respectively, in membrane patches containing a few hundred channels. The presence of the γ1 subunit allows the visualization of distinguishable opening events even at large negative voltages. We determined the product *NPo* using the Poisson distribution (**Eqs. 7-8**, see **Methods**) to fit the area under the peaks of an all-points histogram from long-time single-channel recordings (**Fig. 2C-D-Fig. S3B**). We estimate the number of channels in the membranes using nonstationary noise analysis of the activation currents (**Fig. S3C;** *43, 44*). Once *N* is known, the *Po(V*) data obtained by fitting the theoretical probability density function (*PDF*, **Eq. 8**, see **Methods**) to the experimental *PDF* (**Fig. 2C-D**) allow us to obtain *L*_*0*_ by extrapolation of the *Po(V*) data to 0 mV (**Fig. 2E**). γ1 increases *L*_*0*_ by 92-fold compared to the *L*_*0*_ determined using channels formed by the α subunit alone (**Fig. 2E**, see **Table S1**). The value obtained for *L*_*0*_ of the BK(α) channel is in reasonable agreement with that obtained previously by several groups (*43, 48–51*).

**Figure 2.**
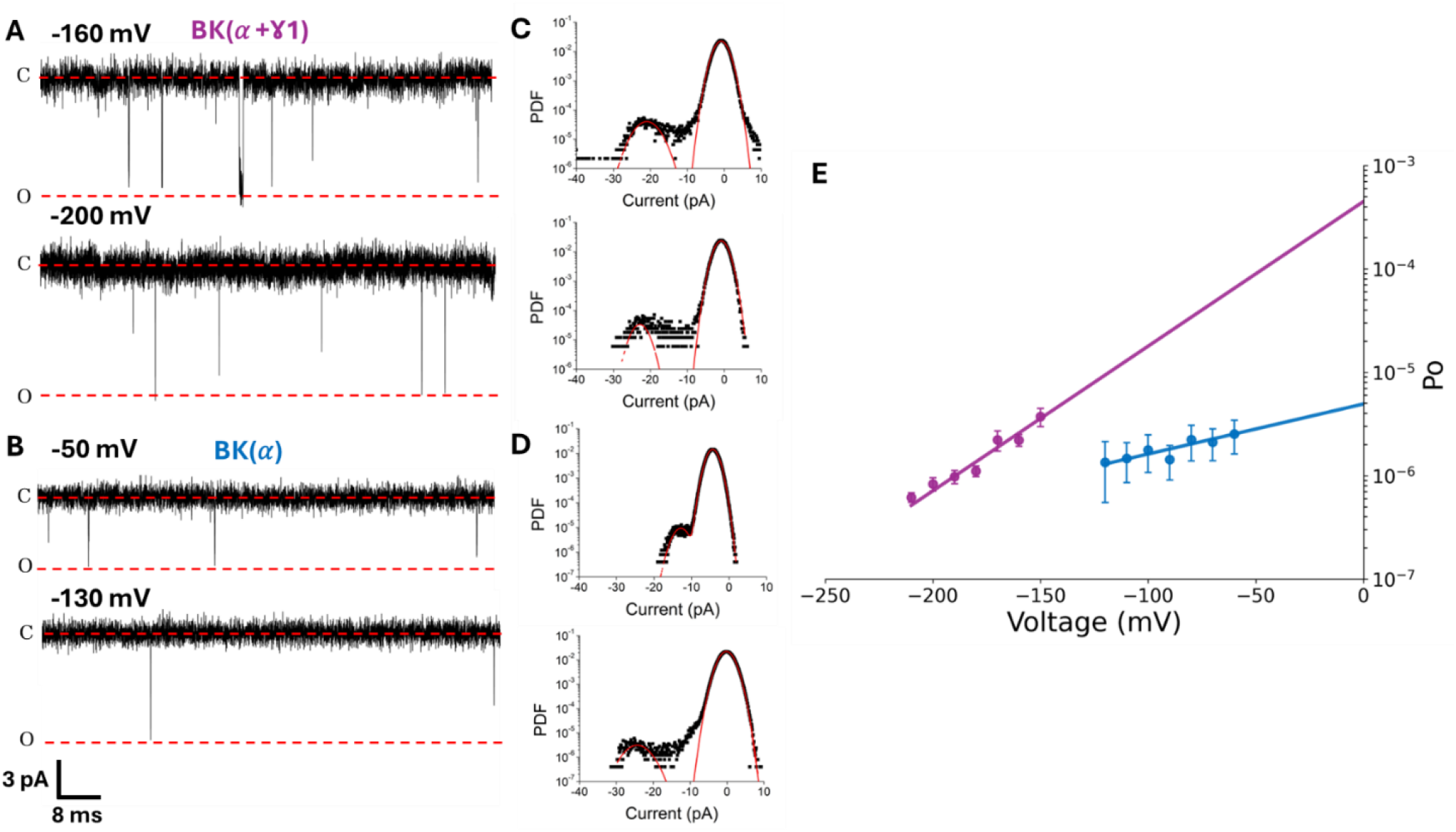
Unitary current analysis of BK in the presence of the γ1 subunit reveals changes in the open-close equilibrium. (**A**) Unitary recordings for hundreds of channels for BK(*α*+γ1) and (**B**) BK(*α*) recorded in inside-out membrane patches at the indicated voltages and under symmetric 110 mM K^+^ conditions and zero μM free internal Ca^2+^. All-point histograms were constructed using 10-20 s of the single-channel recordings on the left, as shown in (**C**) BK(*α*+γ1) and (**D**) BK(*α*). (**E**) Open probabilities at extreme negative voltages for BK(*α*) (blue, N = 5) and BK(*α*) + γ1 (purple, N = 11) were fitted to the straight-line ln(Po) = ln(Lo) + z_L_FV/RT with the following parameters for BK(*α*): L_0_ = 4.9·10^-6^ ± 0.9·10^-6^ y z_L_ = 0.36 ± 0.06 while for BK(*α*+γ1): L_0_ = 4.5·10^-4^ ± 2.9·10^-4^ y z_L_ = 0.74 ± 0.06.

We fitted the *P*_*o*_ data in the whole voltage range to Eq. 3, allowing *D* to vary freely while restricting the value of *J*_*0*_ and *z*_*J*_ to those obtained from fitting the Q(V) data to Eq. 2, and *L* to the values obtained of *L*_*0*_ and *z*_*L*_ extracted from Fig. 2E. Fig. 3A shows that the increase in *L*_*0*_ by the presence of γ1, is accompanied by an increase in the allosteric factor *D* describing the coupling between the voltage sensor and the pore opening from 25.6 to 84.4 (see **Table S1**), which is smaller than the one reported earlier (*D* = 412; (*17)*). Thus, our results show that the leftward shift in the G(V) curve induced by the γ1 is due to the increase in the VSD-PD coupling, accompanied by an appreciable enhancement of *Po* due to the increase in *L*_*0*_. Quantitatively, the reduction in the energy necessary to open the channel is defined as 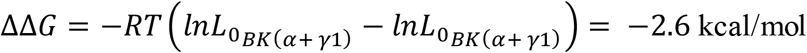. In contrast, the reduction in VSD-PD coupling energy is defined as ΔΔ*G* = −*RT*(*lnD*_*BK*(*α*+ *γ*1)_ − *lnD*_*BK*(*α*)_) = −0.7 kcal/mol per voltage sensor activation. Notably, we found that a 2-fold increase in *z*_*L*_, from 0.36 to 0.74, accompanies the significant rise in *L*_*0*_ due to the presence of γ1 (**Table S1**). Due to the lack of knowledge about the molecular nature of the voltage dependence of the closed-pen reaction, it is difficult to propose a hypothesis about the origin of the increase in *z*_*L*_. However, given that the transmembrane helix of γ1 is kinked and tightly packed against the BK voltage-sensor (*38–40*), the subunit may be stabilizing a BK channel conformation that modifies the electric field where the charges or dipoles that produce the voltage dependence of *L* are displaced.

**Figure 3.**
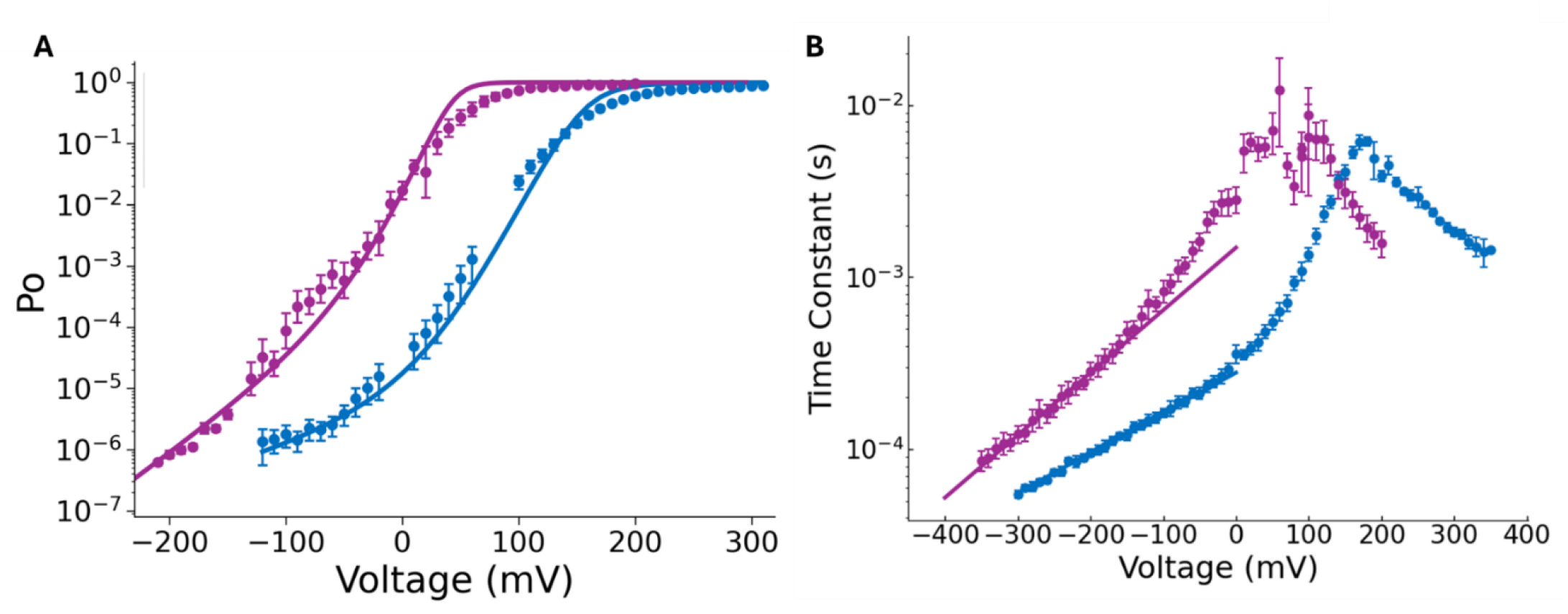
Voltage dependence of the BK(*α*+ γ1) channel open probability and K^+^ current kinetics. (**A**) P_O_ over the whole range of voltages for BK(*α*) (blue, N = 5) and BK(*α*+γ1) (purple, N = 11-7). Data were fitted using Eq. 3, leaving only D as a free parameter, with J and L obtained from the results shown in Figs. 1, 2, and **Table S1**. BK(*α*): D = 25.6 ± 4.9, while BK(*α*+γ1): D = 84.4 ± 6.4. (**B**) Time constant (τ) – voltage data for BK(*α*) (blue, N = 3-4) and BK(*α*+γ1) channels (purple, N = 3-8). At extreme negative voltages, τ was adjusted to the straight line: ln(γ_0_) = ln(γ_0_,_0_) _+_ zγFV/RT for C_0_–O_0_ transition, where γ_0,0_ is the backwards rate at zero mV and z_δ_ the rate voltage dependence. For BK(*α*): γ_0,0_ = 3590 ± 69 (s^-1^) and z_γ_ = 0.136 ± 0.002, while BK(*α*+γ1): γ_0,0_ = 675 ± 41 (s^-1^) and z_γ_ = 0.212 ± 0.005.

In the absence of Ca^2+^ and assuming that the resting-active transition of the voltage sensors is in equilibrium with respect to the closed-open reaction, the time constant-voltage curve, *τ(V*), is given by the relation (kinetics scheme in **Fig. S1B**;49):

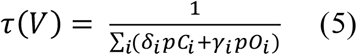

where the subindex *i* = 1, 2, 3, 4 is the number of voltage sensors activated and *δ*_*i*_ and *γ*_i_ are the voltage-dependent forward and backwards rate constants of the individual C_i_-O_i_ transitions, respectively, and *pC*_*i*_ and *pO*_*i*_ are the conditional occupancies of the closed and open states, respectively. Notice that at very negative voltages, the C_0_-O_0_ transition determines τ(V), i.e., *τ(V) = γ*_*0*_^*-1*^. At zero mV, *γ*_0,0_ decreases 5-fold in the presence of the γ1 subunit (see **Table S2**) and the voltage dependence z_γ_, increases from 0.136 for BK(α) to 0.212 for BK(*α*+γ1) suggesting that the γ1 subunit affects the O-C transition explaining the slowing down of the deactivation kinetics and a 29 % of the increase in the voltage dependence of *L* (**Fig. 3B-Fig. S4-S5**). Since *L*_*0*_ = *δ*_*0,0*_*/γ*_*0,0*_, knowing *L*_*0*_ and *γ*_*0,0*_, we can calculate *δ*_*0,0*_. The γ1 subunit increases the forward rate constant *δ*_*0,0*_ by 17-fold compared to the *δ*_*0,0*_ obtained for the BK(*α*) channels (see **Table S2**). This result also corroborates those of Yan and Aldrich (*18*), who found a decrease in *γ*_0,0_ of about 4-fold, albeit with no change in z_γ_. Overall, our results allow us to conclude that, in the absence of Ca^2+^, the main effect of γ1 is to modulate the C-O equilibrium, primarily by destabilizing the closed configuration of the BK channel, thereby facilitating the opening reaction.

### The slow gating charge movement reveals that the γ1 subunit allows BK channel activation from deeper closed states

The kinetic scheme in Fig. S1A predicts that the increase in *D* and *L* induced by the presence of the γ1 should modify the kinetics of the OFF-gating charge (Q_OFF_), as it increases the open-closed equilibrium, allowing the channel to enter open states from deeper closed states. In other words, channels can open with fewer active voltage sensors. Fig. S6A shows OFF-gating currents in response to voltage pulse protocols of increasing duration for the BK(α) channel, designed to characterize their kinetics by integrating the total Q_OFF_. As found previously, as the duration of the depolarizing pulse increases, a slow component develops in the Q_OFF_ vs pulse length in BK(*α*) channels until reaching a constant value (*Q*_*ss*_; **Fig. S6B**; *41, 50*). The *Qss* data obtained using a pulse duration of 25 ms is plotted vs. voltage in Fig. S6C and fitted using Eq. 2 with a *z*_*J*_ = 0.54 and a *V*_*H*_ = 145 mV.

Fig. 4A, shows how a slow component in the gating current records develops as a function of the ON voltage pulse duration in BK(α+γ1) channels (**Fig. 4A**). Fig. 4B shows the time course data of the Q_OFF_ obtained from the gating currents, fitted using a double exponential function at the indicated voltages. Fig. 4C shows the Q_ss_(V) data obtained with a 25-ms pulse duration plotted against voltage. The Q_ss_(V) curve is shifted to the left by 74mV compared to the Q(V) curve obtained by time-integrating the fast component of the ON-gating current (**Fig. 1G**) and approaches the G(V) curve (purple open points; **Fig. 1C**). As a comparison, we show in Fig. 4C the G(V) and Qss(V) curves for the BK(α) channel, noting that the Qss is shifted to the left with respect to the G(V) curve. The results shown in Fig. 4C demonstrate that the γ1 subunit increases the probability that the channels will open at voltages where voltage sensors are not activated by enhancing the closed-open equilibrium.

**Figure 4.**
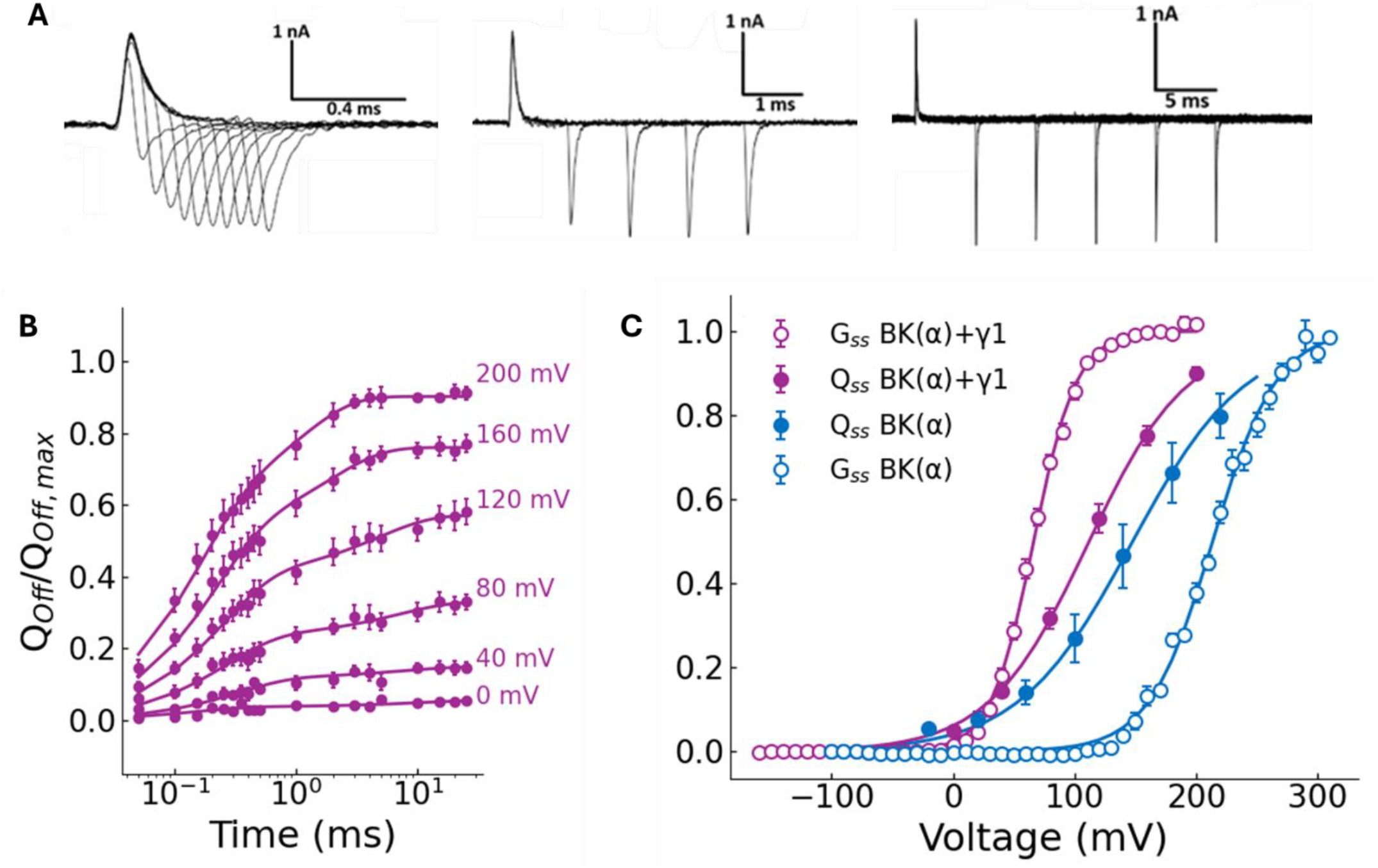
The γ1 subunit enables BK channels to open when only a subset of voltage sensors is active. (**A**) Representative records of gating currents of BK(*α*+γ1) channels, evoked at 200 mV, with different pulse length durations (from 0.05 ms to 25 ms). The total OFF-gating charge (Q_off_) was determined by integrating numerically the OFF-gating current over 0.25 ms. (**B**) BK(*α*+γ1) Q_off_ was normalized by Q_off_,_max_ at 0, 40, 80, 120, 160, and 200 mV, plotted against time (the length for each depolarization pulse; purple filled circle). Data as mean ± SEM, (N = 4), and each dataset was fitted to a biexponential function (see **Methods**) for every voltage. **(C)** Normalized steady state OFF charge (Q_ss_) of BK(*α*+γ1) from experiments like in (**B**) (N = 4) and from BK(*α*) (N = 4, from **Fig. S6**). For comparison, G(V) data from BK(*α*) and BK(*α*+γ1) from Fig.1C were plotted. Q_ss_ were fitted to a two-state model *Q*_*ss*_*/Q*_*ss,-max*_ = 1/[1+exp(−z_Qss_F(V – V_0.5_)/RT)] with the following parameters: V_0.5_ = 147 ± 3 mV and z_Qss_ = 0.54 ± 0.02 for BK(*α*), while a V_0.5_ = 111± 11 mV and z_Qss_ = 0.6 ± 0.1 for BK(*α*+γ1).

### The γ1 subunit does not affect the BK channel Ca^2+^-sensitivity

We next determined the Ca^2+^-dependent activation of BK(α+γ1) channels with robust K^+^ current recordings at multiple Ca^2+^ conditions (**Fig. 5A-D**). Fig. 5E shows a family of G(V) curves obtained from tail currents at various Ca^2+^ concentrations, ranging from 0.01 μM to 1 mM (see **Table S3**). The leftward shift of the G(V) curve (ΔV_0.5_) in the whole Ca^2+^ concentration range is −240 mV, which compares well with that we have obtained previously for the BK channel in the absence of γ1 (shaded zone in **Fig. 5E**; −219 mV (*54*) and −247 mV (*48*)). Corroborating the results from Yan and Aldrich (−210 mV), these findings suggest that γ1 does not affect the BK channel Ca^2+^ activation. However, we can make a more quantitative comparison by fitting Gibbs free energy values (Δ*G*) obtained from the V_0.5_ at different Ca^2+^ concentration data 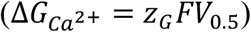 and the relation (*55, 56*):

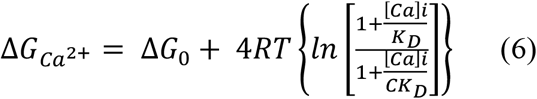

where Δ*G*_0_ is the free energy at zero Ca^2+^, *R* and *T* have their usual meaning, [*Ca*]*i* is the internal Ca^2+^ concentration, *K*_*D*_ the Ca^2+^ binding constant to the closed channel, and *C* is the CTD-PD coupling factor. Eq. 6 assumes one binding affinity (*K*_*D*_) that includes RCK1 and RCK2 binding sites, and that there is no interaction between voltage and calcium sensor, i.e., allosteric factor E = 1. The best fit of the V_0.5_-Ca^2+^ concentration data to Eq. 6 gives no significant differences in *K*_*D*_, and *C* values for BK(α) and BK(α+ γ1) which are very similar to that reported in the literature for the BK(α) channel (41, 45, 51),(*41, 45, 51*), (see **Table S4**). Thus, we confirm previous results that claimed γ1 does not considerable affect the BK channel Ca^2+^ sensitivity (*17*), in opposition to recent findings suggesting a decrease in Ca^2+^ sensitivity and an increase in CTD-PD coupling factor *C* (*39*).

**Figure 5.**
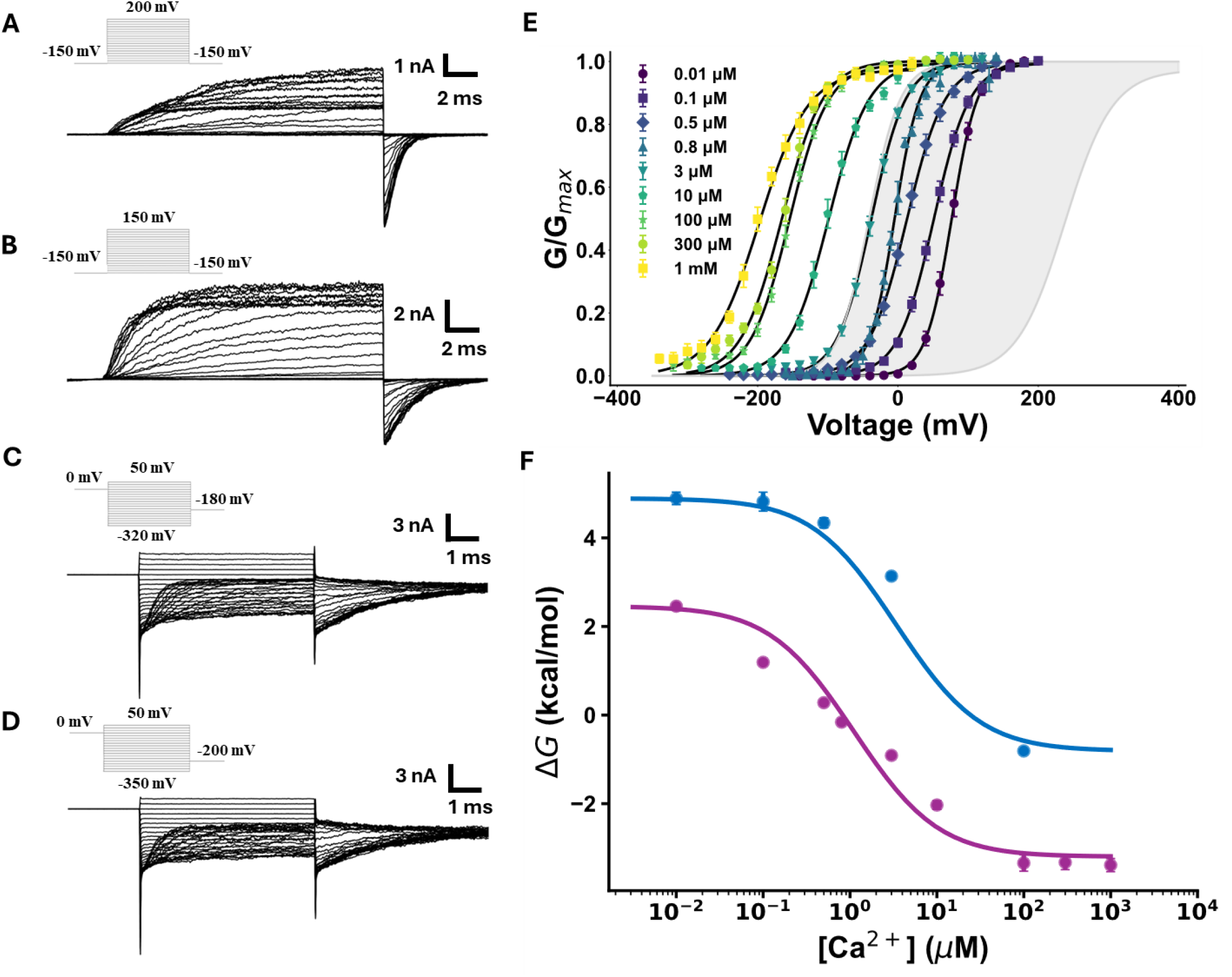
γ1 subunit modulation of Ca^2+^-sensitivity. Representative ionic currents recorded in inside-out configuration under symmetric 110 mM K^+^ conditions and (**A**) 0.01 μM Ca^2+^, (**B**) 0.8 μM Ca^2+^, (**C**) 300 μM Ca^2+^, or (**D**) 1 mM Ca^2+^ for patches containing BK(*α*+γ1) channels. (**E**) Normalized conductance versus voltage plots (G(V)) constructed from tail currents for intracellular Ca^2+^ conditions in the range of 0.01 μM −1 mM were adjusted to a two-state model (see **Table S3**). (**F**) ΔG versus calcium concentration plot for BK(*α*+γ1) channels (purple) and BK(*α*) (blue). The data were fitted using Eq. 6. for BK(*α*+γ1): K_D_ = 0.3 ± 0.2 µM and C = 11.3 ± 1.1 and BK(*α*): K_D_ = 1.0 ± 0.5 µM and C = 11.5 ± 4.1.

## Discussion

### The Horrigan–Aldrich Allosteric Model Provides a Mechanistic Understanding of γ1 Action

Yan & Aldrich (*18*) concluded that γ1 modulates BK channels through a significant increase in the coupling between the VSD and channel opening (allosteric factor *D*; **Fig. S1A**). An increase in *D* from 21 to 412 was enough to explain the leftward shift in the BK voltage activation curve. However, by determining the voltage sensor and pore equilibria, respectively, *J* and *L*, we greatly restricted the degrees of freedom of the fit to the H-A model (only *D* in our case). On the other hand, Redhardt et al. (*39*) proposed a mechanism by which γ1 stabilizes the active state of BK(α) VSD. Moreover, the impact of γ1 on the BK Ca^2+^ sensitivity is under debate. While Yan & Aldrich (*18*) proposed that this was not affected by γ1, Yamanouchi et al. (*40*) results indicate that γ1 weakens the BK(α) channel Ca^2+^ sensitivity, while favoring the active conformation of RCK1 Ca^2+^ binding site. All this evidence points to a controversial subject that has not yet landed into a common answer to the question: how γ1 operates molecularly to modulate the function of BK(α) channels? Motivated by the previous question, we used the Horrigan-Aldrich allosteric gating model. (*18, 43*) to assess the mechanism by which γ1 subunit produces BK channels to be able to activate at negative potentials in the absence of Ca^2+^ (*18, 43*).

Notably, our results show that the closed-open equilibrium is greatly affected by γ1, which increases *L*_*0*_ by more than 90-fold and interestingly, augments *z*_*L*_, accounting for BK channel voltage-dependent activation regardless of VSD activation. Increasing *L*_*0*_ orders of magnitude have been described previously in point mutants of BK (*57*), however in these mutants the voltage dependence remained unaltered. This effect, produced by the γ1 subunit suggests a novel mechanism of action involving the modification of the electric field presumably around the pore domain. The increase in *L*_*0*_ is partially due to a 5-fold decrease in the closing rate constant (*γ*_*0*_) but mainly caused by a 17-fold increase in the forward rate constant (*δ*_*0*_) of the closed-open reaction. Our results and the fitting of the P_o_(V) data using the allosteric model indicate that the increase in *D* is smaller than that obtained by Yan and Aldrich (*18*), and are not consistent with their proposal that *L* was unaffected by the γ1 subunit. Still, their results show a 4-fold decrease in *δ*_*0*_, an observation not discussed in their study. Regardless of these differences, in our results, the effect of the γ1 on the allosteric factor *D* is at a lesser scale, increasing from 26 in the BK(α) to 84 in BK(*α*+γ1) channels. However, even if we consider that D = 400 in BK(*α*+γ1) as reported by Yan and Aldrich (*18*), the reduction in VSD-PD coupling energy per voltage sensor activation (1.7 kcal/mol) is still smaller than the reduction in the energy necessary to open the channel promoted by the γ1 subunit (2.8 kcal/mol).

### The resting-active equilibrium of the VSD is not affected by the γ1 subunit

The present results appear to be incompatible with the mechanism proposed by Redhardt et al. (*39*), who suggested that the positive charges (arginines) contained in γ1 C-terminal stabilize the VSD active configuration through long-range electrostatic interactions. An electrostatic bias imposed by the internal arginine cluster of γ1 should affect the resting and active equilibrium of the voltage sensors, thereby shifting the Q(V) curves towards negative voltages. The fact that the gating charge activation curve is not affected by γ1 when the channels are closed discards the possibility that a long-range electrostatic effect is involved in the mode of action of γ1. Moreover, the fact that γ1 does not affect the resting-active equilibrium between closed states does not support the proposal that γ1 increases the probability of opening at negative potential by locking the active configuration of the VSD through the conformational changes in the S3 transmembrane segment (*40*). In Fig. 1, we demonstrate that γ1 does not stabilize the VSD active configuration when the channels are closed and that the BK channel reaches its maximum open probability when fewer than 20% of the voltage sensors are in their active configuration. In other words, the data indicate that γ1 increases *D* to stabilize the VSD active configuration when the channels are open. The HA allosteric gating model gives a parsimonious explanation of these results. By increasing the equilibrium constant that defines the closed-open equilibrium (*L*) and the coupling between the voltage sensor and pore opening (*D*), γ1 alters the primary activation pathway (see **Fig. S1B**).

### Ca2+ sensitivity and physiological implications of the BK(*α*+γ1) complex

Regarding Ca^2+^-dependent activation, Yan & Aldrich (*18*) previously suggested that γ1 does not modify the BK channel Ca^2+^ sensitivity. However, Yamanouchi et al. (*40*) found a decrease in BK channel Ca^2+^ sensitivity induced by γ1 and suggested that the interaction between the RCK1 domain and the γ1 arginine cluster promoted and increase in the RCK-PD coupling factor *C*. Nonetheless, our experiments at multiple Ca^2+^ concentrations, showing a slight increase in Ca^2+^ affinity and no alteration in coupling factor *C*, indicate that there is no effect in CTD-PD coupling nor Ca^2+^ affinity decrease due to γ1 subunit. The increase in Ca^2+^ affinity is nothing but a result of an energetic bias that the γ1 subunit imposes in the voltage-dependent activation curves, which is more noticeable when calculating and comparing the difference in Δ***G*** between saturating Ca^2+^ and free Ca^2+^ conditions (ΔΔ***G*;** see **Table S4**), which is the same in the presence of the γ1 subunit. Therefore, the γ1 subunit positions BK(α+γ1) within the activation window at resting intracellular Ca2+, providing a mechanistic basis for robust physiological K^+^ fluxes (see **Fig. 5E**).

### Towards a structural and mechanistic understanding of γ1 modulation of the BK channel

Our data are in line with Kallure et al.(*38*) proposal based on the BK(α+γ1) structure, which posits that the γ1 transmembrane domain kink promotes a stabilization of the voltage sensor domain helices (S0-S3), which favors the active configuration of S4 by primarily affecting VSD-PD communication (*38*). However, the main departure of previous interpretations of the modulatory effects of the γ1, is that γ1 mainly increases the channel open probability by destabilizing the closed state and not by increasing the allosteric coupling factor *D*. This kink may also be involved in the increase of PD voltage-dependence *z*_*L*_, most likely reducing the electric field fraction that charges must traverse to open the channel.

Molecular modeling (*58–60*) and structural studies (*61, 62*) demonstrate that the open-closed transition implies a rotation and an approximation of the S6 transmembrane helices, making the BK channel’s internal vestibule more hydrophobic and decreasing the pore radius. These structural characteristics of the closed conformation led to the hydrophobic gating hypothesis (*58–60*). The closed BK channel structure also shows the appearance of membrane-facing fenestrations (*61–64*). Molecular simulations indicate that lipids can bind to the membrane-facing fenestrations and access the inner pore cavity, thereby increasing its hydrophobicity (*65*). The γ1 subunit transmembrane domain kink may destabilize the closed configuration by introducing a “brake” to the configurational changes that lead to the closed state. However, to answer this question, a well-resolved structure of the closed BK(*α*+γ1) channel and molecular modeling will allow us to compare the stability of the closed channel configuration with that of the BK(*α*) channel.

## Materials and Methods

### Molecular Biology and Channel Expression

*Xenopus laevis* oocytes were utilized as the heterologous expression system for BK(α) and BK(α+γ1) expression. The α subunit of the human BK channel (GenBank: U11058) was gift of L. Toro from the University of California, Los Angeles. The cDNA for hSlo1 was subcloned into the pBSTA vector. The mutant 2D2A (D362A/D367A) for the RCK1 Ca2+-binding site (Xia et al., 2002) and the 5D5A mutant for the RCK2 Ca2+-binding site, also known as the calcium bowl (D894A/D895A/D896A/D897A/D898A; (*66*), were kindly provided by M. Holmgren from the National Institutes of Health in Bethesda, MD. Dr. Jiusheng Yan from the University of Texas MD Anderson Cancer Center in Houston, TX, and Dr. Richard Aldrich from the University of Texas at Austin, TX provided the cDNA coding for the human γ1 subunit (LRRC26). The cRNA was checked and sequenced. The restriction enzyme NotI was used to digest the cDNA. The linearized DNA was transcribed using the mMESSAGE mMACHINE T7 transcription kit (Ambion, Austin, TX, USA). The RNA obtained was quantified using a Nanodrop, which measured the 260/280 and 260/230 ratios to determine purity, and agarose gel electrophoresis to assess quality. Finally, Xenopus laevis oocytes were injected with 50 ng of cRNA for BK(α), and when injected with γ1 in a ratio of [RNAγ1]:[RNAα] = 5, they were incubated in an ND96 solution (in mM: 96 NaCl, 2 KCl, 1.8 CaCl_2_, 1 MgCl_2_, 5 HEPES, pH 7.4) at 18°C for 1–4 days before electrophysiological recordings.

### Electrophysiology

All recordings were conducted using the patch-clamp technique in the inside-out configuration. For K^+^ ionic currents (I_K_), we used symmetrical solutions composed of mM: K-MeS 110 (KOH 108 and KCl 2), HEPES 10, and 5 EGTA for the ‘ free’ Ca2+’ solution (~0.8 nM, based on the presence of ~10 μM contaminant [Ca2+] (*67*), adjusted to pH 7.4). For test solutions at different Ca2+ concentrations (0.03 – 1000 μM), CaCl_2_ was added to achieve the desired free [Ca2+], and 5 mM EGTA (0.1–0.5 μM) or HEDTA (0.8–10 μM) was used as a calcium buffer. No Ca2+ chelator was used for ≥100 µM Ca^2^+ solutions, and the free Ca^2^+ concentration was estimated using the WinMaxChelator Software and verified with a Ca^2^+ electrode (Hanna Instruments). All experiments were performed at room temperature (20–22°C). Potassium currents at different Ca^2^+ concentrations in the same oocyte were obtained by excising the patch and washing it with an appropriate internal solution using at least 10 times the chamber volume. Pipettes for macroscopic and gating currents were fabricated using borosilicate capillary glass (1B150F-4, World Precision Instruments, Sarasota, FL, USA) and pulled using a horizontal puller (Sutter Instrument) and fire-polished (Narishige, London, UK). The pipettes for macro patches were designed to achieve a pipette resistance in the range of 0.5–1.5 MΩ. Data were acquired using an Axopatch 200B (Molecular Devices). Voltage commands and current outputs were filtered at 20 kHz using an 8-pole Bessel low-pass (Frequency Devices). For current recordings, we sampled using a 16-bit A/D converter (Digidata 1550B; Molecular Devices), sampling at a rate of 250 kHz.

For gating currents (I_g_), a solution containing (mM): X-MeS 110, HEPES 10, and EGTA 1, adjusted to pH 7.4, where X = N-methyl-D-glucamine (NMDG) in the bath solution was used. To avoid ionic currents through BK(α+γ1) channels were blocked using 2 µM paxilline. In the pipette solution X = tetraethylammonium (TEA). The voltage and current were filtered analogically using an 8-pole Bessel low-pass filter (Frequency Devices, Ottawa, IL, USA) at 10 kHz, through a 16-bit analog-to-digital converter (Digidata 1550B, Axon Instruments) with a sampling frequency of 500 kHz. The gating currents were measured using giant patches with 0.5– <1 MΩ pipettes. In the case of BK(α+γ1) channels the gating charge was estimated by fitting the gating current time course using of two exponentials of which only the fast component was considered when calculating the gating charge.

### Determination of the open probability (*P*_*o*_)

For *Po* determination, macro-patches containing hundreds of channels were used. Tail current analysis was used to determine *Po* values > 0.01. For Po < 0.01 values, membranes with many channels and measurements of single-channel currents at large hyperpolarization were used. Nonstationary noise analysis (*46, 47*) was used to calculate the number of channels (*N*) in the patch and the unitary current (*i*). The average current and variance of isochrones were obtained from 200 current relaxation records in response to a depolarizing voltage pulse or during the tail after depolarization. After channel counting, membrane current was acquired for 5–30 s at Vm values ranging from 60 to −220 mV, voltages where the instantaneous current corresponds to a few open channels. To determine the open channel probability (*NPo*), we generated all-point histograms of the number of times a given current, *I*, was observed. The ordinate values of the histogram were divided by the integral over the current axis to obtain an experimental probability density function, *PDF*. The *PDF* is modeled as a sum of 10 normal Gaussian functions (**Eq.7**) weighted by coefficients, *Pk*, obtained from Poisson’s distribution (**Eq. 8**) where the expected number of open channels is *NPo*

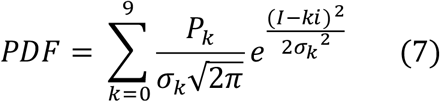

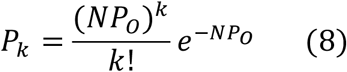

where *P*_*k*_ is the probability of having *k* open channels. The mean value of each Gaussian is the amplitude of *k* unitary currents, *i*, while the noise of one open channel defines the corresponding variance:

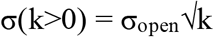

For recordings showing two or more simultaneous open channels, the variance of the closed channel, σ(k=0), was empirically found to be 0.75 times the variance of the open channel, σ_open_. This restriction was applied to fit the model, where only single channels were detected. We also adjusted the unitary current, *i*, to resemble the instantaneous I/V relationship from tail currents in deactivation protocols. At extremely negative potentials, where single-channel openings are very brief, the acquisition filter distorts the shape of normal histograms between Gaussian peaks (*68*). Because in this condition, the model PDF lies below the experimental data points between the two Gaussian peaks, we corrected the *NPo* as the area under the *k=1* Gaussian, plus the area between the model and experimental PDFs divided by 2. By previously subtracting the baseline, the free parameters of the Poisson-Gauss fit are *NPo* and σ_open_.

## Funding

FONDECYT Regular #1230265 (R.L.)

ANID Millennium Science Initiative Program P09-022F (R.L.)

National Institute Award RO1GM030376 (R.L.)

PhD ANID fellowship #21202097 (F.E.)

## Author contributions

Conceptualization: R.L.

Methodology: R.L., O.A., J.P.C., F.E., A.P-P., M.F., W.C-U.

Investigation: F.E., A.P-P., M.F., W.C-U.

. Formal Analysis: R.L., O.A., J.P.C., F.E., A.P-P., M.F.

Supervision: R.L.

Writing—original draft: R.L.

Writing—review & editing: R.L., O.A., J.P.C., F.E., A.P-P., M.F., W.C-U.

## Data and materials availability

All data are available in the main text or the supplementary materials.

## Notes

### Competing Interest Statement

The authors have declared no competing interest.

