## Supplemental Materials for "Tuning the gate and the gear: The LRRC26 (γ1) subunit modulates intrinsic gating and voltage-sensor coupling of the BK channel"

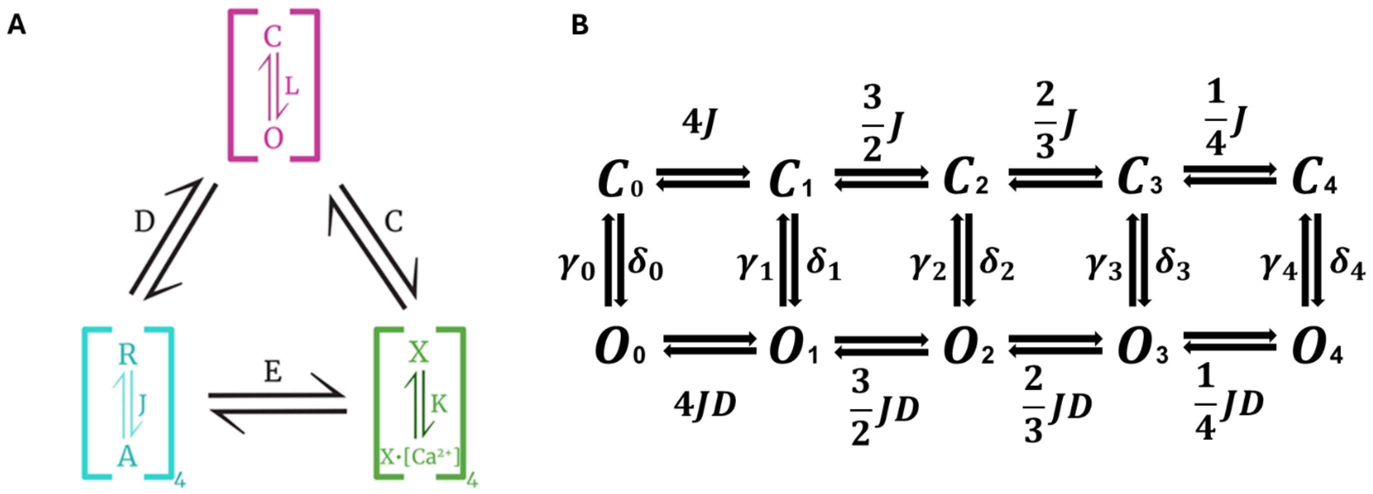


**Supplementary Figure 1. HA allosteric gating model**. (**A**) Scheme of HA allosteric model (*42*) indicates the possible conformations of the PD (C-O), VSD (R-A), and $Ca^{2+}$ sensors (X-X·[$Ca^{2+}$]) in each of four α-subunits. The equilibrium constant, L, defines channel opening; The equilibrium constant J defines VSD activation; the binding constant K determines $Ca^{2+}$ binding. The allosteric relationships between the modules are described by the allosteric factors D, C, and E. (**B**) Gating kinetics scheme. In this scheme, the horizontal $C_i$-$C_i$ and $O_i$-$O_i$ transitions are assumed to be in equilibrium for the $C_i$-$O_i$ transitions, where $\delta_i$ and $\gamma_i$ are the forward and backward rate constants defining the $C_i$–$O_i$ transitions.

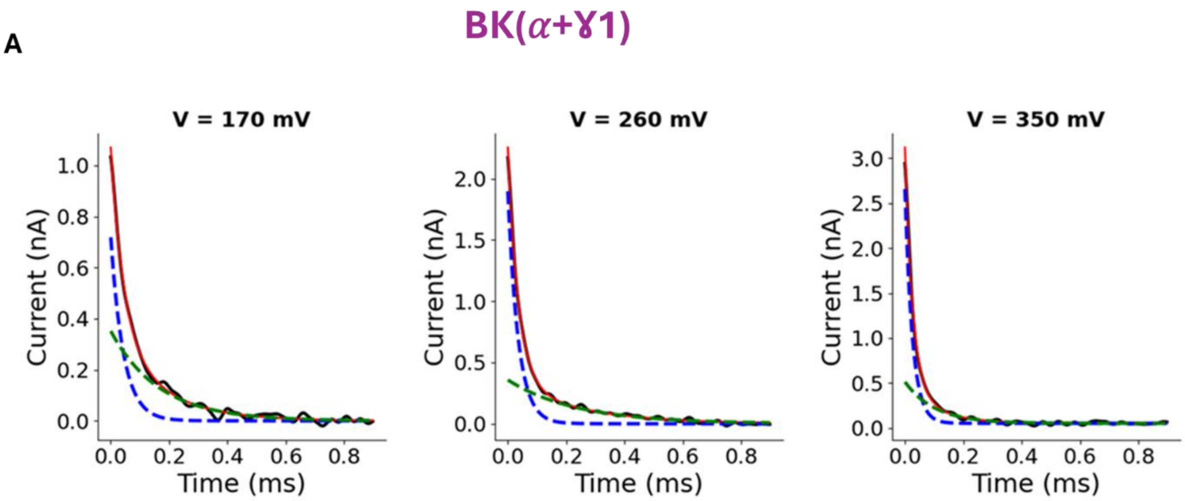


**Supplementary Figure 2. Double exponential analysis of ON gating currents from BK(α+γ1) channels.** (**A**) Representative biexponential fit (red) to gating currents (black) at three depolarizing pulses: 170 mV (left), 260 mV (center), and 350 mV (right). The function used for the biexponential fit was $I_g=A_{fast}e^{-t/\tau fast} + A_{slow}e^{-t/\tau slow}$ where A is the amplitude of each exponential. The gating current time constant (τ) for the fast (blue) and slow (green) component, respectively, resulted for each voltage was: 170 mV: τ-fast = 44 ± 2 µs; τ-slow = 167 ± 9 µs, 260 mV: τ-fast = 35.4 ± 0.4 µs; τ-slow = 237 ± 9 µs, and 350 mV: τ-fast = 25.5 ± 0.4 µs; τ-slow = 103 ± 6 µs.

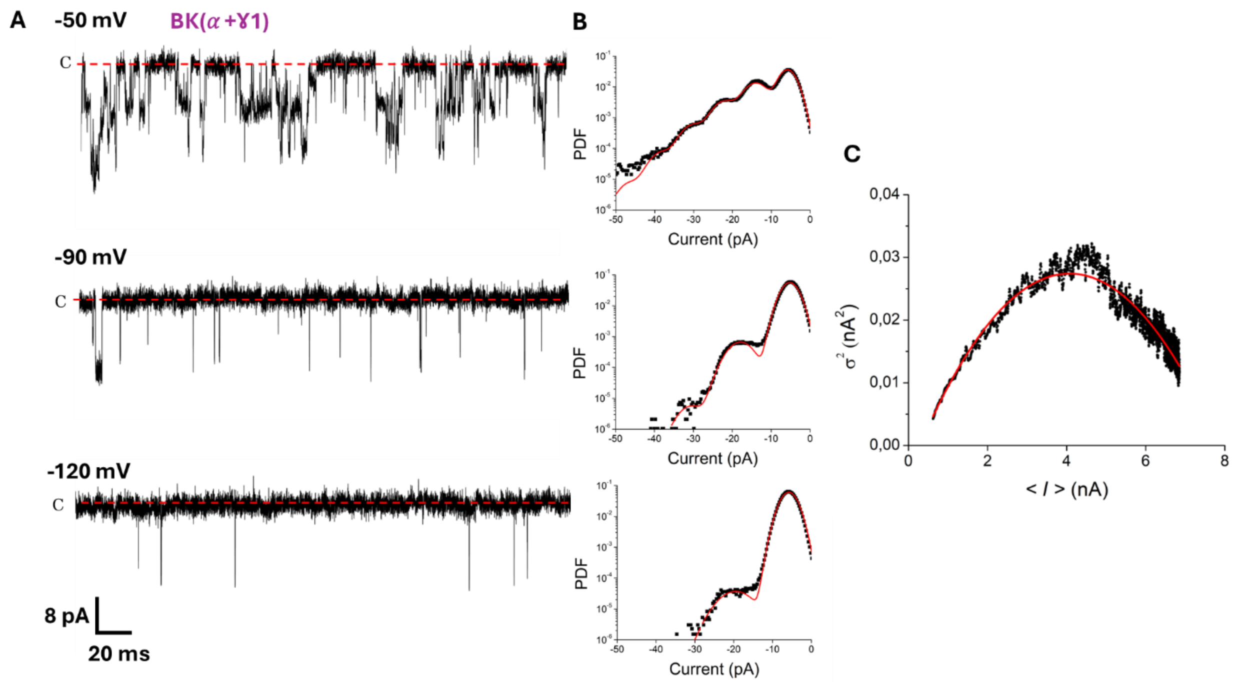


**Supplementary Figure 3. Poisson distribution analysis for BK(α+γ1) unitary currents.** (**A**) Unitary recordings for hundreds of channels of BK(α+γ1) were recorded in inside-out membrane patches at the indicated voltages and under symmetric 110 mM $K^+$ conditions and zero free internal $Ca^{2+}$. All-point histograms of the opening events were constructed using 10-20 s of the single-channel recordings on the left, as shown in (**B**), and fitted to a Poisson distribution to calculate $NP_O$ at multiple voltages and construct $P_O$-V curves in the whole voltage range. **(C)** The number of channels (N) was obtained using non-stationary noise analysis on activation currents for each recording. In this example, the variance ($\sigma^2$)-mean current (<*I*>) data was fitted (red) using the relation: $\sigma^2 = i\langle I\rangle - \langle I\rangle^2/N$, where $i$ is the single-channel current with $i$ = 20 pA and $N$ = 304 channels. <*I*> was obtained from the average of 200 current records elicited using a 110-mV voltage pulse.

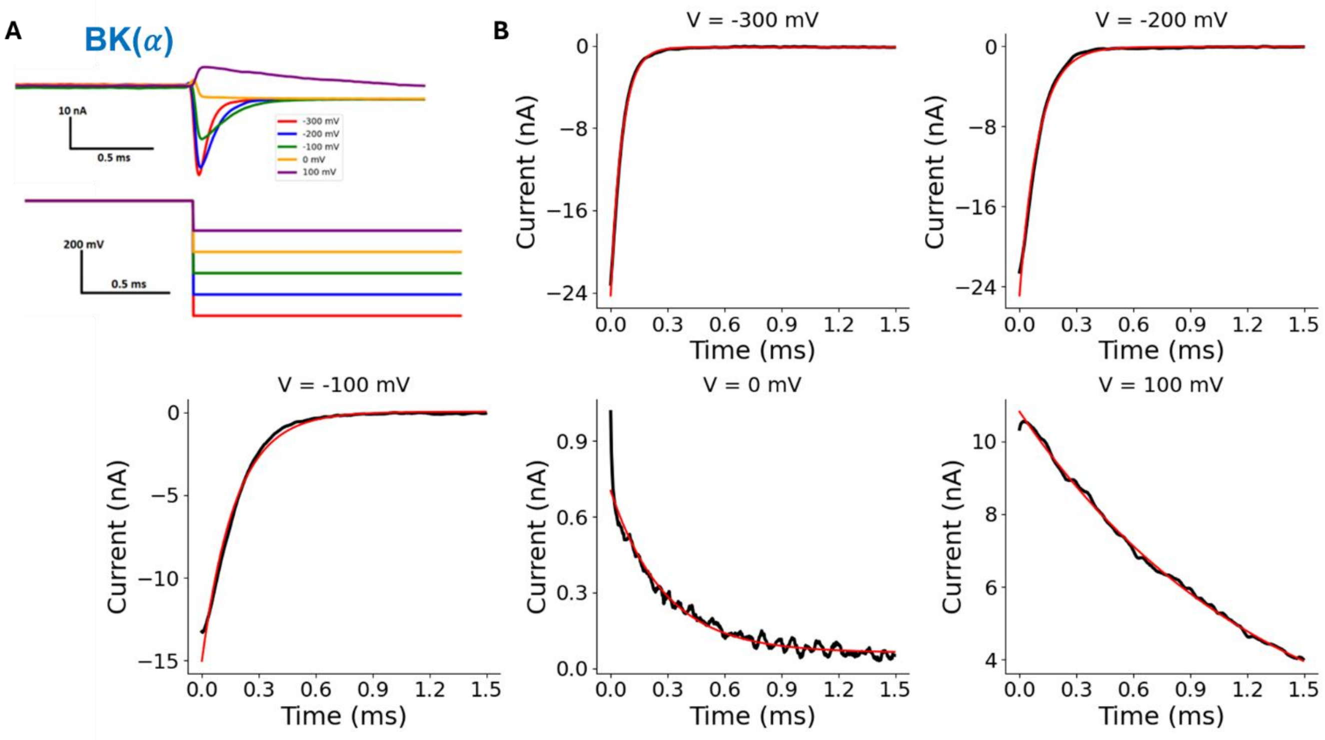


**Supplementary Figure 4. Monoexponential analysis of deactivation kinetics from BK(α) channels.** (**A**) Representative currents (top panel) of ionic currents from BK($\alpha$) channels, evoked in response to an instantaneous voltage protocol (bottom panel) at different hyperpolarization potentials (see the inset for color symbology). (**B**) Representative monoexponential fits (in red) to tail currents (in black) from (**A**). Time constants of deactivation were estimated for five different potentials: 59.3 ± 0.2 μs (-300 mV), 99.8 ± 0.7 μs (-200 mV), 175.6 ± 1.5 μs (-100 mV), 285± 6 μs (0 mV), and 1310 ± 20 μs (100 mV).

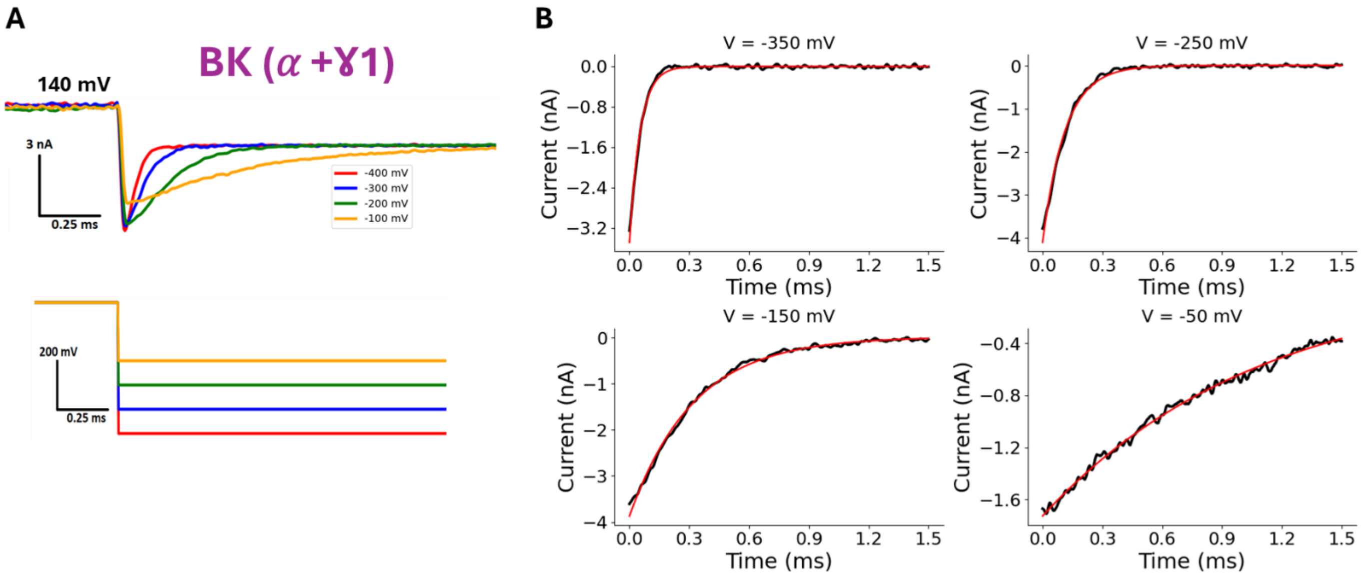


**Supplementary Figure 5. Analysis of deactivation kinetics from BK(α+ γ1) channels.** (**A**) Representative currents (top panel) of ionic currents from BK(α+γ1) channels, evoked in response to an instantaneous voltage protocol (bottom panel) at different hyperpolarization potentials (see inset for color symbology). (**B**) Representative monoexponential fits (in red) to tail currents (in black) from (**A**). Time constants of deactivation were estimated for five different potentials: 53.4 ± 0.4 µs (-350 mV), 113.3 ± 0.8 µs (-250 mV), 320.6 ± 2.3 µs (-150 mV), and 1091.2 ± 233 µs (-50 mV)

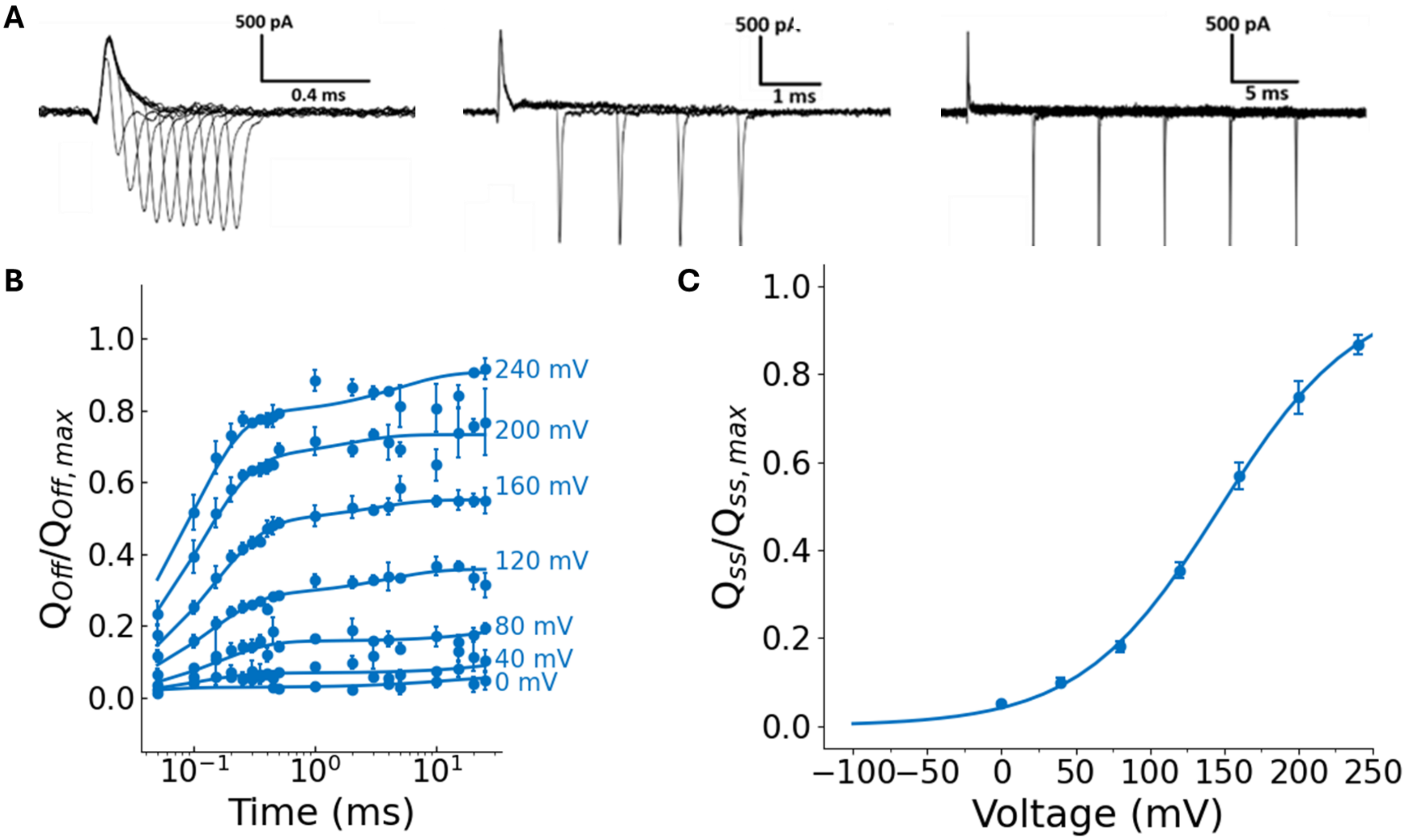


**Supplementary Figure 6. BK($\alpha$) OFF-gating charge kinetics.** (**A**) Representative records of gating currents of BK($\alpha$) channels, evoked at 200 mV, with different pulse length durations (from 0.05 ms to 25 ms). The total OFF-charge ($Q_{off}$) was determined by numerically integrating the OFF-gating current during 0.25 ms. (**B**) $Q_{off}$ was normalized by $Q_{off,max}$ at 0, 40, 80, 120, 160, 200, and 240 mV, plotted against time (the length for each depolarization pulse; mean ± SEM, N = 4), and each dataset was fitted to a biexponential function (see **Methods**) for every voltage. (**C**) Normalized steady state OFF-charge ($Q_{ss}$) of BK($\alpha$) from experiments like in (**B**) (N = 4). $Q_{ss}$ were fitted to a two-state model $Q_{ss}/Q_{ss,-max}$ = 1/[1+exp(-$z_{Qss}$F(V–V0.5)/RT)] with the following parameters: $V_{0.5}$ = 146.9 ± 2.8 mV and $z_{Qss}$ = 0.544 ± 0.019.

| Parameter | BK($\alpha$ +γ1) | BK($\alpha$) |
|---|---|---|
| $V_{0.5}$ (mV) | 80.9 ± 0.3 | 218.3 ± 0.8 |
| $z_G$ | 1.37 ± 0.02 | 0.85 ± 0.02 |
| $V_{H,}$ (mV) | 185 ± 1.5 | 178 ± 0.4 |
| $z_Q$ | 0.58 ± 0.04 | 0.59 ± 0.04 |
| $J_0$ | 0.013 ± 0.001 | 0.029 ± 0.005 |
| $z_J$ | 0.58 ± 0.04 | 0.59 ± 0.04 |
| $L_0$ | $4.5*10^{-4} \pm 2.9*10^{-4}$ | $4.9*10^{-6} \pm 0.9*10^{-6}$ |
| $z_L$ | 0.74 ± 0.06 | 0.36 ± 0.06 |
| $D$ | 84.4 ± 6.4 | 25.6 ± 4.9 |

**Supplementary Table 1.** Summary of BK($\alpha$) and BK($\alpha$+γ1). $V_{0.5,}$ and $V_H$ correspond to the half-activation voltages for G(V) and Q(V) curves (**Fig. 1**), respectively. The other parameters correspond to the HA model.

| Parameter | BK($\alpha$ +$\gamma$1) | BK($\alpha$) |
|---|---|---|
| $\gamma_{0,0}$ ($s^{-1}$) | 675.0 ± 41.1 | 3587.2 ± 68.8 |
| $z_{\gamma}$ | 0.212 ± 0.005 | 0.136 ± 0.002 |
| *$\delta_{0,0}$ ($s^{-1}$) | 0.3 ± 0.2 | 0.017 ± 0.004 |
| *$z_{\delta}$ | -0.53 ± 0.06 | -0.22 ± 0.06 |

**Supplementary Table 2.** Summary of BK($\alpha$+$\gamma$1) and BK($\alpha$) kinetic. Values (*) were calculated from $\gamma_{0,0}$ and $L_0$.

| $[Ca^{2+}]_{Free}$ (μM) | BK($\alpha$ +ɣ1) (mV) | BK($\alpha$) (mV) |
|---|---|---|
| 0.01 | 80.9 ± 0.3 (9) | 218.3 ± 0.8 (8) |
| 0.1 | 55.9 ± 0.3 (8) | 233.5 ± 8.5 (4) |
| 0.5 | 12.7 ± 0.6 (5) | 193.9 ± 1.7 (4) |
| 0.8 | -4.8 ± 0.8 (3) | |
| 3 | -37.8 ± 0.8 (5) | 126.6 ± 1.3 (4) |
| 10 | -98.8 ± 1.5 (3) | |
| 100 | -154.7 ± 1.7 (7) | -35.6 ± 1.0 (4) |
| 300 | -165.2 ± 1.7 (4) | |
| 1 (mM) | -197.7 ± 1.8 (7) | |

**Supplementary Table 3.** Summary of BK($\alpha$+γ1) and BK($\alpha$) half activation voltages for a two-state model from the family of G(V)s at intracellular $Ca^{2+}$ concentrations in the range of 0.01 μM -1 mM, followed by the number of replicas in parentheses.

| Parameter | BK($\alpha$ +ɣ1) | BK($\alpha$) |
|---|---|---|
| $K_D$ (μM) | 0.3 ± 0.2 | 1.0 ± 0.5 |
| $C$ | 11.3 ± 1.1 | 11.5 ± 4.1 |
| $*\,\Delta\Delta G(0.01\ -100\ uM)$ (kcal/mol) | 5.68 ± 0.04 | 5.9 ± 0.2 |

**Supplementary Table 4.** Summary of BK($\alpha$+γ1) and BK($\alpha$) $Ca^{2+}$ dose-response parameters.
